# Longitudinal Proteomic Profiling of T Cell Differentiation In Vivo Reveals Biochemical Remodeling Underlying Exhaustion

**DOI:** 10.1101/2024.05.14.593504

**Authors:** Christian M. Beusch, Abdelhameed S. Dawood, Ahmet Ozdilek, Sarah Welbourn, Youssef M. Zohdy, Christopher M. Monaco, Alexandra S. Flegle, Ahmed Mahgoub, Sakshi Malik, Christina Niavi, Akil Akhtar, Carly Roman, Autumn A. Gavora, David E. Gordon, Mohamed S. Abdel-Hakeem

## Abstract

CD8 T cell exhaustion impedes immune responses to cancer and chronic infections, and a biochemical understanding of exhaustion is essential to improving immunotherapy. Here, we present the first longitudinal protein abundance and phosphoproteomic analysis of antigen-specific CD8 T cells undergoing differentiation in vivo during acute (LCMV-Armstrong) and chronic (LCMV-Clone 13) infection. Comparing protein abundance across the two infection conditions identified over 180 known and novel exhaustion-associated proteins, including proteins missed by transcriptional analyses. Phosphoproteomic analysis identified >900 differentially regulated phosphosites on >400 proteins, including known inhibitory phosphosites on PD1, PAG1, SHP-1/PTPN6, SLAMF1/CD150. We also calculated phosphosite conservation across mammals, to direct follow-up studies towards sites with likely essential function. Lastly, our analysis uncovers exhaustion-associated kinases with clinical-stage inhibitors, underscoring the translational utility of our dataset to guide immunotherapy development. Together, our datasets define a biochemical atlas of T cell exhaustion in vivo, shedding light on the molecular mechanisms of T cell dysfunction.

## Introduction

CD8 T cells are central in combating tumors and intracellular pathogens. During acute infection, naïve T cells are activated, giving rise to effector cells (T_EFF_) which differentiate into memory T cells (T_MEM_) upon clearing the pathogen. This is accompanied by extensive changes in the phenotype and cellular profiles of T cells^1–3^. However, under chronic infections and cancer, antigen persistence leads to alternative T cell differentiation through the “exhaustion” pathway, resulting in exhausted T cells (T_EX_) that progressively lose function and exhibit distinct phenotypic, transcriptional, and epigenetic profiles compared to T_EFF_/T_MEM_ cells ^4–9^. The biochemical landscape of antigen-specific T cell differentiation *ex vivo* is understudied, limiting our ability to therapeutically modulate T_EX_ cells.

While transcriptomic and epigenetic approaches offer valuable insights into gene expression and regulation, they do not directly measure the functional molecules driving cellular processes – the proteome. Several studies have reported discrepancies between RNA and protein levels, highlighting the importance of direct protein measurements^10–12^. More importantly, analysis of post-translational modifications (PTMs) of proteins, e.g., phosphorylation, offers biochemical insights that are not accessible to transcriptional readouts. Notably, the central importance of phosphorylation for T cell activation, and its antagonism by inhibitory signaling pathways, are central to the regulation of T cell response to antigen stimulus^13–15^. Therefore, analyses of protein abundance and phosphorylation in the context of T cell differentiation afford an enhanced perspective on the mechanisms underlying T cell differentiation and exhaustion. Previous proteomic studies of T cell differentiation have primarily been conducted *in vitro*, often utilizing non-specific stimulation and focusing on the early stages of T cell activation^16–18^. Other investigations have profiled immune cells’ proteome in the context of viral infections or cancer, demonstrating that proteomic analyses can uncover key proteins overlooked in transcriptomic datasets—some of which hold promise as immunotherapeutic targets^19, 20^. However, comprehensive proteomic analysis of *ex vivo* CD8 T cells in an antigen-specific model across T cell differentiation states has not been performed.

Here, we profiled the dynamics of proteomic alterations during the differentiation of antigen-specific CD8 T cells *in vivo* in the context of acute versus chronic infection with lymphocytic choriomeningitis virus (LCMV) Armstrong (Arm) and Clone-13 (CL13) strains, respectively. Overall, our comprehensive analysis of global proteomics aligns with established knowledge of T cell biology, highlighting key canonical inhibitory receptors, metabolic pathways, and transcription factors associated with the different states of T cell differentiation. Crucially, our analysis reveals inconsistencies between mRNA and protein abundance for several differentially expressed proteins, underscoring the indispensable role of proteomics in capturing the true biochemical landscape of T cell differentiation. Additionally, our phosphoproteomic analysis highlights known phosphorylation sites mediating inhibitory signaling, and also identifies a multitude of novel phosphorylation sites associated with T cell differentiation that merit further study. Using kinase substrate enrichment analysis, we identified state-specific kinases, e.g., T_EX_-specific kinases, as targets for kinase inhibitor interventions. Notably, we applied a sequence conservation-based prioritization approach to focus on phosphorylation sites of potential functional importance and highlight several understudied conserved phosphorylation sites identified in our dataset. Given the widespread adoption and translational relevance of the LCMV mouse model, our dataset provides a powerful resource for deeper insights into T cell differentiation and serves as a valuable complement to other omics approaches.

## Results

### Longitudinal proteomic profiling of CD8 T cells reveals dynamic proteome modulations

For decades, the LCMV mouse model has been instrumental in the phenotypic, transcriptional, and epigenetic characterization of T cell differentiation. A central strength of this system is the ability to compare mice infected with the Clone 13 strain (LCMV-CL13), which induces chronic infection, to those infected with the Armstrong strain (LCMV-Arm), which induces acute infection. This comparison specifically reveals exhaustion-associated molecular programs in T cells during chronic infection, offering valuable insights into the underlying mechanisms of exhaustion that revealed successful immunotherapeutic approaches^1, 5, 8, 21–23^.

Leveraging the power of the LCMV model, we employed liquid chromatography-tandem mass spectrometry (LC-MS/MS) to profile proteomic alterations of transgenic LCMV-GP33-specific P14 CD8 T cells at different stages of chronic and acute infection (Fig.1A). P14 cells were adoptively transferred into congenically distinct recipient mice, which were either infected with chronic LCMV-CL13 or acute LCMV-Arm, respectively generating exhausted T cells (T_EX_) or effector/memory T cells (T_EFF_/T_MEM_). Mice were sacrificed, and P14 cells were isolated and sort-purified across a time course at 8, 15, and 30 days post-infection (hereafter d8, d15, and d30), representing early, intermediate, and late stages of infection^1, 5, 8, 21, 23^. Naïve P14 cells were sorted as a benchmark for protein expression in the naïve state. Our flow cytometry analyses reproduced established phenotypes of T_EX_ and T_EFF_/T_MEM_ (Fig.1B-C, Extended Data Fig.1). Sorted P14 cells were processed for global protein abundance analysis by LC-MS/MS. We quantified over 6,600 proteins across the entire dataset with a robust correlation in reproducibility across all biological replicates, and quantified proteins within a broad dynamic expression range (Extended Data Fig.2A-E). We confirmed concordance between canonical T cell differentiation markers’ expression measured by flow cytometry and mass spectrometry-based proteomics (Fig. 1D, Extended Data Fig.2F-G), including markers associated with activation and effector phenotype (KLRG1, CX3CR1, CD44, and granzyme B) as well as exhaustion-associated markers (PD1, Tox, and CD39/*Entpd1*). Overall, differentiation of CD8 T cells during acute and chronic infection led to significant alterations in their proteome, delineated by two distinct trajectories (Fig.1E), confirming unique differentiation of T_EX_ compared to the T_EFF_/T_MEM_ on the proteome level as previously shown phenotypically^22, 23^, transcriptionally^8, 21^, and epigenetically^1, 5^. Quantitatively, activation from the naïve T cell state to either T_EFF_ or early-T_EX_ (naïve vs. d8) was accompanied by changes in the abundance of ∼1,000 proteins, which is significantly higher than modulations occurring at further stages of differentiation (Fig. 1F). For acute infection, significant changes continued as the cells transitioned from early T_EFF_-dominated response (d8) towards a T_MEM_-dominated response (d15), but to a much lesser extent from intermediate to late T_MEM_. Crucially, in the context of chronic/CL13 infection, proteome modulation persisted throughout the early-to-intermediate exhaustion transition and the intermediate-to-late exhaustion transition (Fig.1F). Overall, global protein alterations quantified across our time course analysis of acute and chronic LCMV infections align with previously reported transcriptional and epigenetic studies^5, 8, 21^.

**Figure 1:**
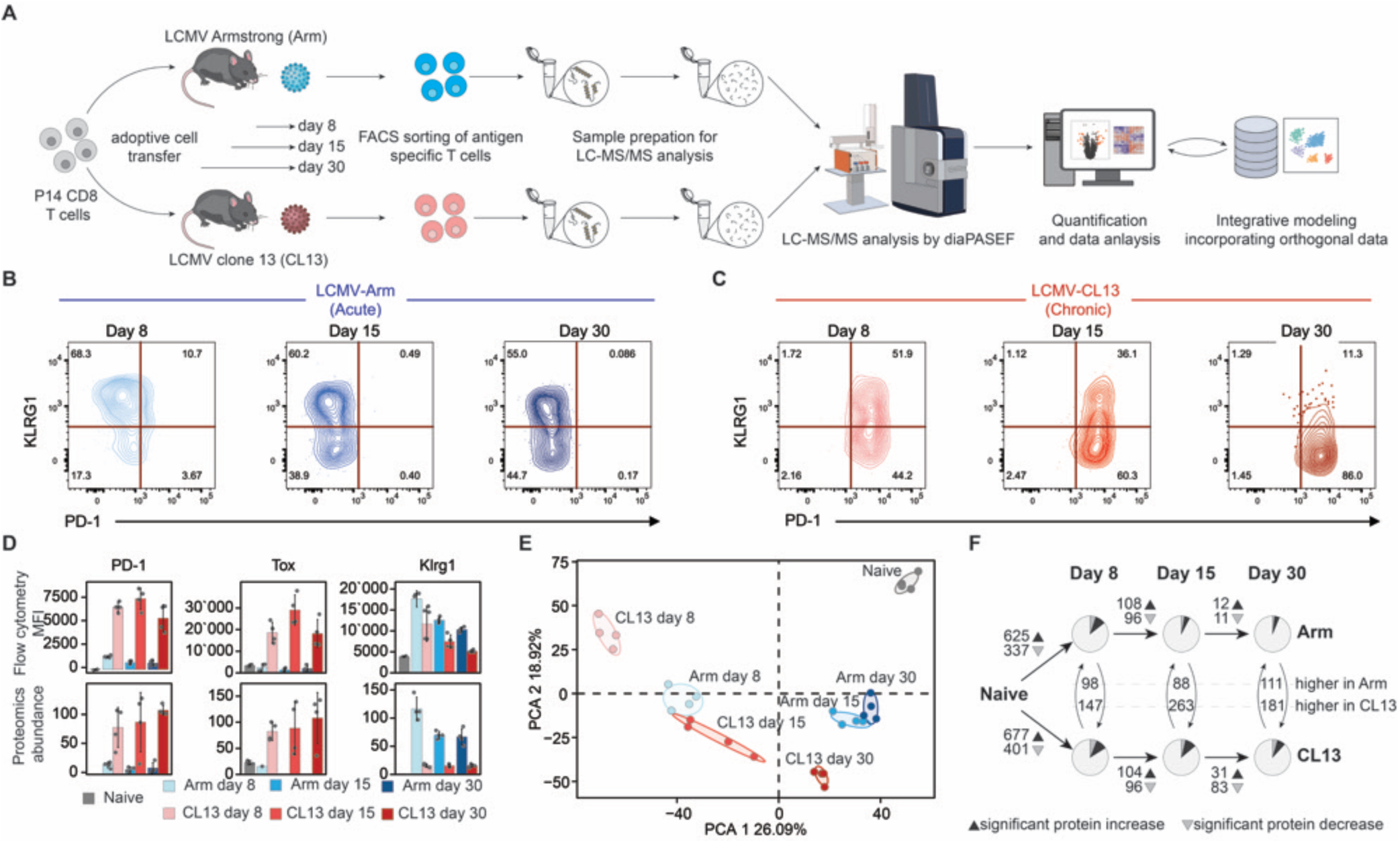
Longitudinal proteomic profiling of antigen-specific CD8 T cells distinguishes proteome alterations under acute and chronic infection conditions. **(A)** Overview of the experimental design to analyze proteome-wide changes in CD8 T cells differentiating during acute (Armstrong) and chronic (CL13) LCMV infections. **(B, C)** Representative flow cytometry dot plots showing expression levels of the exhaustion marker PD-1 and activation marker KLRG1 on transgenic GP33-specific P14 CD8 T cells. **(D)** Comparative analysis of protein expression per-cell (MFI) assessed by flow cytometry (top) and mass spectrometry-based proteomics (bottom). N = 4, with error bars representing mean ± standard deviation. Naïve B6 mice were used as flow cytometry staining controls. The absence of a measurement bar indicates proteins not quantified by mass spectrometry. **(E)** Principal component analysis (PCA) of proteomics samples, highlighting distinct clustering of acute versus chronic infection conditions. **(F)** Quantification of protein abundance changes across the T cell differentiation trajectory in acute (top) and chronic (bottom) LCMV infection. Dark wedges indicate upregulated proteins (log₂FC > 1), while grey wedges represent downregulated proteins (log₂FC < - 1), p < 0.05, one-sided ANOVA. Curved lines denote the number of differentially expressed proteins comparing acute/Armstrong versus chronic/CL13 at each post-infection time point.

### Delineating T cell proteome alterations during acute versus chronic infection identifies exhaustion-specific proteins

T cell exhaustion is a hallmark of cancer and chronic viral infection, representing a divergent differentiation path for T_EX_ distinct from T_EFF_/T_MEM_^1, 6–8, 21, 23–31^. We harnessed the full potential of the LCMV mouse model, which enables comparison of antigen-specific CD8 T cells with the same specificity during both acute and chronic infection, to dissect the proteome dynamics accompanying differentiation of T_EFF_/T_MEM_ and T_EX_.

We applied unsupervised hierarchical clustering to group proteome alterations of T_EFF_/T_MEM_ and T_EX_ T cells into distinct clusters based on their longitudinal dynamics (Fig.2A, Extended Data Fig.3A-B), followed by gene ontology (GO) enrichment analysis to functionally categorize proteins within each cluster (Extended Data Fig.3C). For example, T_EX_-enriched modules reflected in cluster 4 included Eomes and Lag3, cluster 6 included PD1, cluster 8 included Tox, and cluster 11 included Tigit and CD244/2B4. These clusters were associated with GO terms including negative regulation of cell activation and defense response (cluster 4 and cluster 6); inflammatory response and negative regulation of virus replication (cluster 8); and regulation of cell killing and positive regulation of cytokine production (cluster 11). Focusing on specific protein families highlighted several biological processes, including downregulation of respiratory chain complex 1 and mitochondrial translation-associated proteins during chronic infection (Fig.2B and Extended Data Fig.3D), features of T_EX_ associated with metabolic dysfunction and impaired oxidative phosphorylation^32, 33^. We also observe a spike in protein expression of MCM complex associated with cell proliferation in T_EFF_ cells, which returns to baseline in T_MEM_, but is sustained in T_EX_ (Fig. 2B, bottom panel) – this is consistent with previous literature^23^.

**Figure 2:**
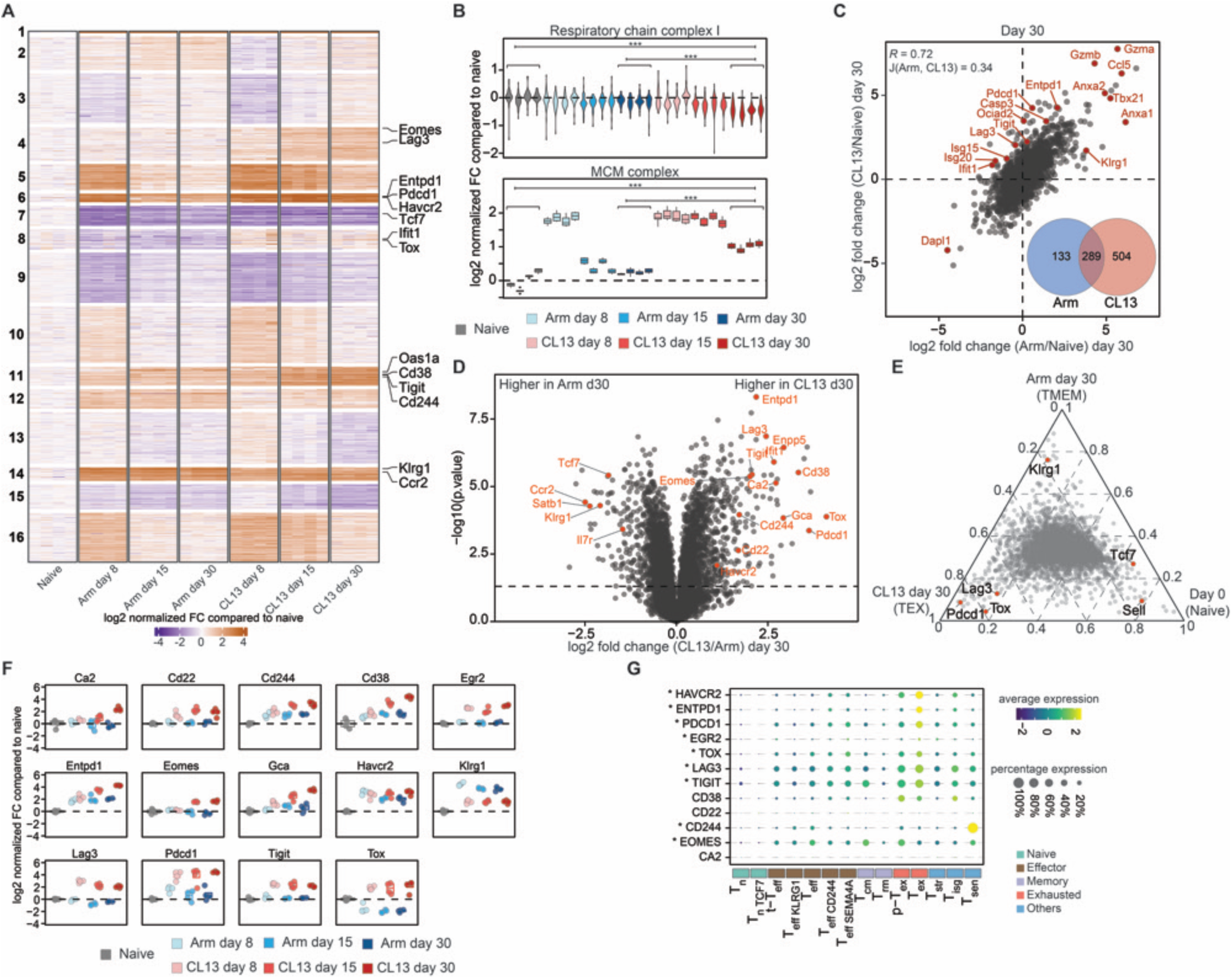
Exhaustion-specific dynamic changes of the T cell proteome in the context of chronic LCMV-CL13 infection. (**A**) Unsupervised hierarchical clustering of differentially expressed proteins across acute (Arm) and chronic (CL13) infections, highlighting protein clusters with distinct temporal expression patterns. (**B**) Distribution of protein expression changes for components of the respiratory chain complex I and minichromosome maintenance (MCM) complex across all samples, illustrating metabolic and cell cycle regulation dynamics. (**C**) Scatter plot comparing protein expression in memory and exhausted T cells at day 30 post-infection relative to naïve T cells. The Venn diagram quantifies the overlap of significantly regulated proteins between the two conditions. Sample correlation was determined using Pearson correlation, while the Jaccard index assessed the similarity of significantly regulated proteins. (**D**) Volcano plot displaying differential protein abundance in P14 CD8 T cells from Arm- versus CL13-infected mice at day 30 post-infection. (**E**) Ternary plot illustrating protein-specific expression patterns across naïve, memory (Arm day 30), and exhausted (CL13 day 30) T cells, with labeled proteins serving as reference points. (**F**) Boxplots of selected proteins from (D), depicting individual proteomics measurements along the T cell differentiation trajectory. (**G**) Bubble plot showing gene expression of selected proteins from (F) across tumor-infiltrating CD8 T lymphocyte (TIL) subsets, based on data from the Pan-Cancer T Cell Atlas^1^. Asterisks denote canonical exhaustion-associated genes. N = 4 for all proteomics samples.

By leveraging time-matched samples of T_EFF_/T_MEM_ from acutely infected (LCMV-Arm) mice and T_EX_ from chronically infected (LCMV-CL13) mice, we can directly compare the distinct differentiation trajectories, analogous to previous transcriptomic and epigenetic studies. Indeed, analyzing LCMV-Arm and LCMV-CL13 in parallel distinguishes the core program of T cell differentiation at the intersection between Arm and CL13, versus context-specific programs unique to either acute or chronic conditions at the early, intermediate, and late stages of infection, respectively (Fig.2C-E, Supplementary Table 1, Extended Data Fig.3E-I). These comparisons confirmed elevated expression of T_EX_ inhibitory surface proteins: PD1/*Pdcd1*, TIM3/*Havcr2*, 2B4/*Cd244*, CD39/*Entpd1*, Lag3, and TIGIT; in addition to the main transcription factor orchestrating the exhaustion program, Tox (Fig.2D). Conversely, several proteins consistently exhibited higher abundance in T_EFF_/T_MEM_, including the activation marker KLRG1, the memory-associated markers CD127/*Il7r* and TCF1/*Tcf7* at the intermediate and late stages, and SATB1 which recruits the NuRD deacetylase complex to repress PD1/*Pdcd1* expression^34^. To identify proteins specifically regulated during T cell exhaustion, we used trinary plot analysis comparing naïve, exhausted (CL13 d30), and memory (Arm d30) T cells. Naïve markers like Sell (CD62L) and Tcf7 were enriched in naïve cells, while exhaustion markers such as Pdcd1, Lag3, and Tox were specific to exhausted T cells (Fig.2E).

While the exhaustion program in T_EX_ is best understood when contrasted with T_EFF_/T_MEM_, there is also value in analyzing the two trajectories on their own. Our longitudinal profiling of protein abundance in T_EFF_/T_MEM_ during acute viral infection revealed dynamic changes throughout transitions from naïve, to effector, to memory (Extended Data Fig.4), and corroborates transcriptional and epigenetic trends previously reported^2, 3, 8, 21^. Studies of *in vitro* T cell stimulation demonstrated extensive proteome remodeling upon activation^12, 16, 18^, however, our study details a comprehensive profiling of the *in vivo* trajectory of T_EFF_/T_MEM_. Our analysis of the T_EX_ proteomic trajectory during chronic viral infection also represents the first insight into the longitudinal *in-vivo* protein abundance dynamics of T_EX_ (Extended Data Fig.5).

### Tumor-associated transcriptomics highlight commonalities of T_EX_ biology across chronic diseases

To validate that our proteomic results are not LCMV-context-dependent, we compared our protein abundance results to the pan-cancer atlas single-cell transcriptomics dataset of human tumor-infiltrating lymphocytes^43^. Across several T_EX_-specific proteins identified in our dataset, we quantified the proportion of cells expressing each transcript across all CD8 T cell clusters (Fig.2G circle diameter), and the average RNA expression (Fig.2G color bar, yellow = highest). Across both canonical markers (*) and less characterized proteins, we consistently observed higher percent expression and greater RNA expression in T_EX_-clusters, demonstrating that T_EX_-specific proteins identified by our proteomic analyses could be extended to the T cell behavior present in human cancer. In several cases, we noted T_EX_-upregulated proteins in our dataset were near/below the limit of detection in the pan-cancer atlas T cell scRNAseq dataset (Fig.2G).

### Integrating our protein abundance and transcriptomics data identified additional proteins implicated in T cell exhaustion

Transcriptional analyses are pivotal to our understanding of T cell differentiation^1, 8, 21^. However, previous studies underscored the partial correlation between mRNA and protein abundance upon T cell activation *in vitro*, highlighting the significance of post-transcriptional mechanisms governing protein expression^10–12, 17, 44^.

To compare transcriptome and proteome changes during T cell differentiation, we performed RNA-seq on the same samples used for proteomics, and compared RNA and protein abundance (Fig.3, Extended Data Fig.6, Supplementary Table 3). This analysis revealed a Pearson correlation coefficient of 0.51 at 30 dpi and 0.32-0.33 at earlier time points (Fig.3A-C), aligning with other published comparisons of RNA vs protein abundance in diverse contexts^10, 19, 45^. Gene ontology analysis of genes with concordant changes in both RNA and protein levels reveals enrichment for functions related to lymphocyte activation and regulation, and cell-cell adhesion. In contrast, genes with discordant RNA and protein changes are enriched for functions such as cell migration, organelle disassembly, RNA capping, and signal peptide processing (Fig.3D–F).

**Figure 3:**
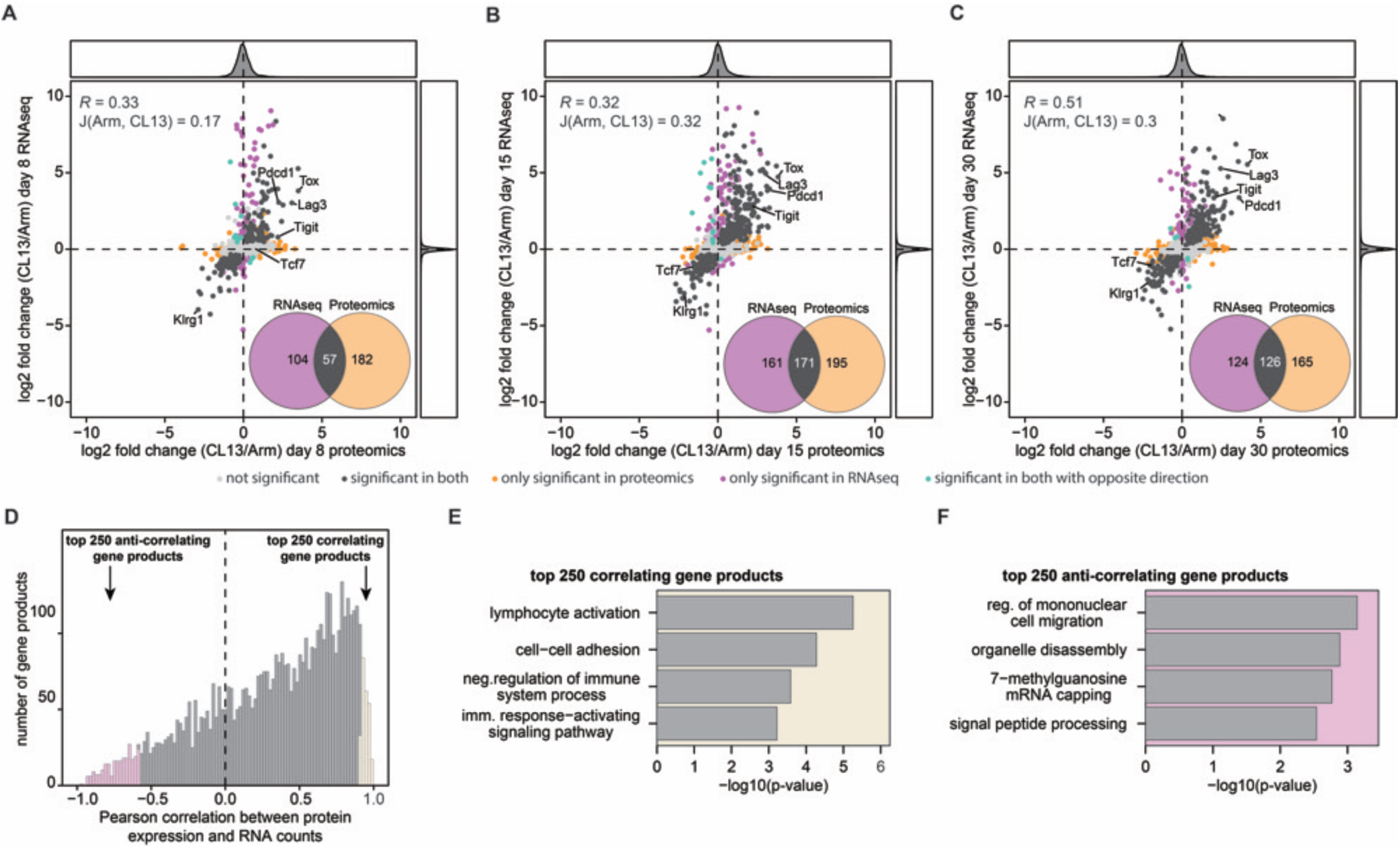
Integration of T cell transcriptional and proteomic profiles reveals variability in transcript-to-protein correlation. (**A–C**) Scatter plots comparing protein expression (proteomics) to mRNA levels (bulk RNAseq) at different time points post-infection. Venn diagrams highlight the overlap of significantly regulated gene products between the two methods. Pearson correlation was used to assess overall sample correlation, while the Jaccard index was used to quantify the similarity of significantly regulated proteins. (**D**) Histogram showing the distribution of Pearson correlation coefficients between protein abundance and corresponding mRNA levels, illustrating the variability in transcript-to-protein relationships. (**E, F**) Gene Ontology (GO) enrichment analysis of the top 250 positively correlated (concordant) and negatively correlated (discordant) gene products. N = 4 for all proteomics samples, N = 3–4 for RNA sequencing.

Generally, we observe concordance between RNA and protein abundance, notably with canonical exhaustion markers including PD1/*Pdcd1*, Lag3, and Tox, which were initially identified via transcriptomics (Fig.3A-C). Together, these analyses validate the use of global proteomics to identify T_EX_-associated markers and uncover novel proteins involved in exhaustion biology, highlighting the critical importance of integrating proteomic and transcriptomic data for a more comprehensive and unbiased understanding of T cell exhaustion.

### Phosphoproteomic analyses reveal exhaustion-specific signaling modularity

Post-translational modifications (PTMs) such as phosphorylation regulate biochemical interactions and signaling pathways underpinning cellular functions^13, 49–51^. Previous studies have explored phosphoproteomic analysis of T cell activation *in vitro*^52, 53^, however, comprehensive phosphoproteomic analysis of antigen-specific T cells directly *ex vivo* has not yet been performed. We performed phosphopeptide enrichment on the same samples as used for our global proteomics analysis (Fig.4A), identifying over 10,000 localized phosphosites with high quantitative overlap and reproducibility between biological replicates (Extended Data Fig.7A-B). Importantly, to derive unbiased biochemical insights, we corrected for protein abundance differences of our phosphoproteomics quantities (Extended Data Fig.7C). Principle component analysis shows clear separation by infection condition and time point post-infection indicating distinct phosphorylation states between sample groups (Fig.4B). Phosphoproteomic analysis uncovered extensive remodeling of signaling networks during the transition from naïve T cells to either T_EFF_ or early-T_EX_ states (Fig. 4C), and dynamic changes persisted throughout T_EX_ differentiation, whereas under acute infection phosphoproteomic changes are reduced as cells transition to T_MEM_ throughout d15 and d30. These findings reinforce the notion that T cell exhaustion is not merely a consequence of phenotypic, epigenetic, transcriptional, or global proteomic changes but is also shaped by distinct post-translational modifications and their impacts on cellular biochemical networks and signaling pathways.

**Figure 4:**
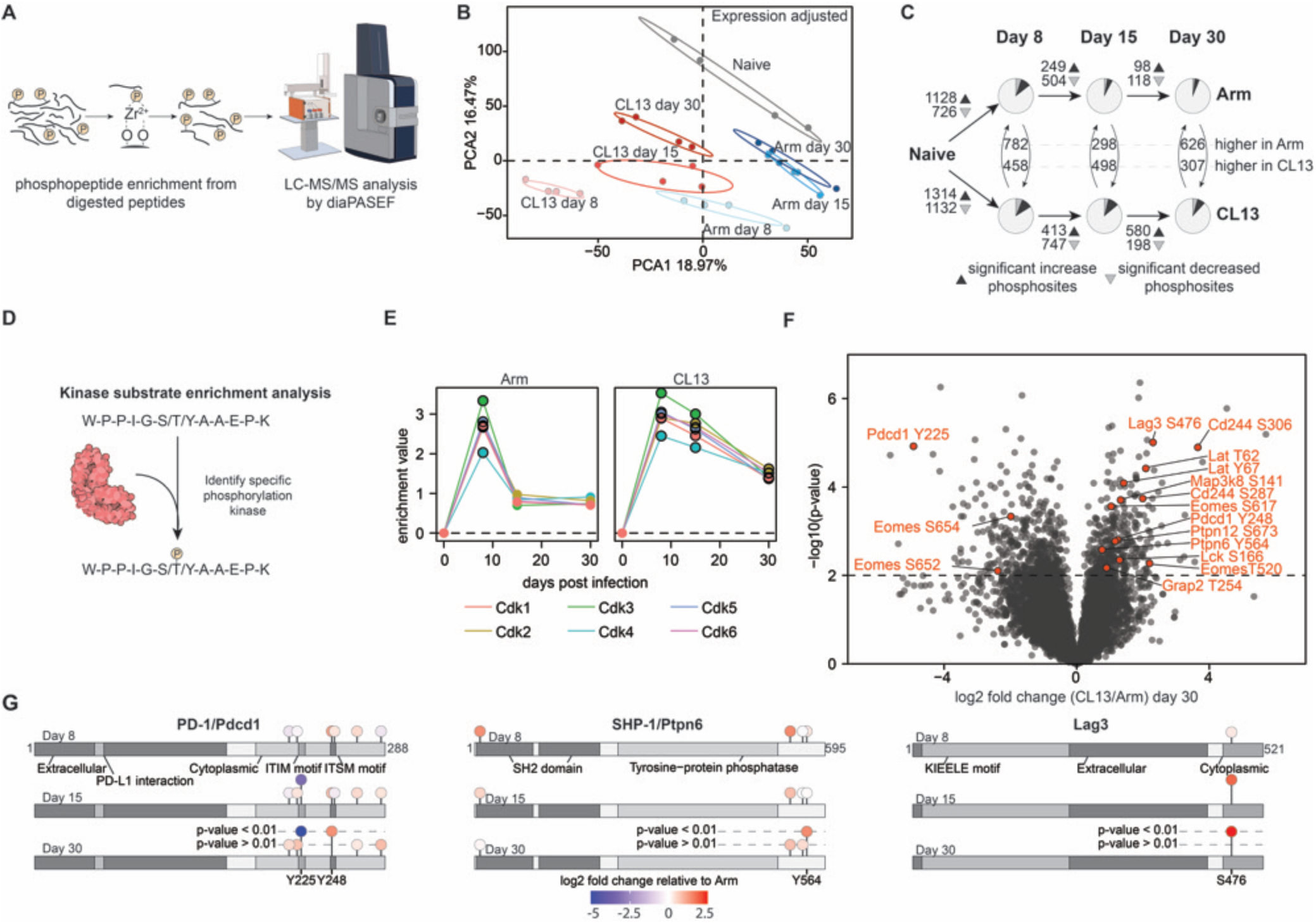
Phosphoproteomic profiling reveals unique kinase activity and phosphorylation sites in the chronic LCMV-CL13 infection context. (**A**) Schematic representation of the phosphopeptide enrichment workflow used for phosphoproteomic data acquisition. (**B**) Principal component analysis (PCA) of phosphoproteomics samples, showing distinct clustering between acute and chronic infection conditions. (**C**) Quantification of phosphosite abundance changes across the T cell differentiation trajectory in acute and chronic viral infections. Dark wedges indicate upregulated phosphosites (log₂FC > 0.75), while grey wedges denote downregulated phosphosites (log₂FC < -0.75), p < 0.01, one-sided ANOVA. (**D**) Overview of Kinase-Substrate Enrichment Analysis (KSEA), a computational method to identify kinases potentially responsible for phosphorylating specific phosphosites. (**E**) Longitudinal kinase activity profiling of selected Cyclin-Dependent Kinases (CDKs) in acute and chronic infection. (**F**) Volcano plot depicting differentially regulated phosphosites in CD8 T cells from Arm- vs. CL13-infected mice at day 30 post-infection. (**G**) Schematic representation of PD-1/Pdcd1 (left), SHP-1/Ptpn6 (middle), and Lag3 (right), highlighting all quantified phosphorylation sites. N = 4 for all proteomics samples.

### Phosphoproteomics reveals kinase signatures of T cell exhaustion

Kinase-substrate enrichment analysis (KSEA) enables inference of upstream kinases responsible for specific phosphorylation events, and thus identifies potential kinase targets for therapeutic intervention^54, 55^ (Fig.4D). Comparing phosphosite changes in the different cell states throughout the differentiation trajectory revealed a consistent trend across cyclin-dependent kinases (CDKs). CDK activity surged at d8 in both conditions, persisting in T_EX_ while returning to quiescent levels in T_MEM_ (Fig.4E, Extended Data Fig.7D, Supplementary Table 4). In late-T_EX,_ CDK activity ultimately diminished, aligning with reduced proliferative capacity in late-T_EX_^23^. Reassuringly, this temporal pattern mirrored the protein abundance trend of MCM complex proteins (Fig. 2B, bottom panel), which are also associated with cell proliferation.

Thus, comprehensive KSEA of our phosphoproteomic data might revealed functionally relevant kinases that orchestrate T cell regulation, pinpointing key regulatory nodes that could serve as therapeutic targets.

### Comparative Phosphoproteomics of Exhausted and Memory T Cells Uncovers Regulatory Mechanisms

Phosphorylation at specific protein residues acts as a molecular switch, regulating protein activity, stability, and localization – and ultimately influencing cell fate and function^52, 63^. In some cases, specific phosphorylation events can initiate or alter entire signaling pathways^64, 65^. With this in mind, we focused on differentially regulated phosphorylation sites between T_EX_ (CL13-d30) and T_MEM_ (Arm-d30), as these sites could provide biochemical mechanistic and therapeutic insights (Fig.4G, Supplementary Table 5). Among the most significantly modulated sites on T_EX_, we identified several located on key inhibitory receptors and transcription factors. These include PD1 Tyrosine 225 (phosphosite notion Y225) and Y248; Lag3 S476; CD244 S287 and S306; and Eomes T520, S617, S652 and S654. Additionally, we observed dysregulated phosphorylation of proteins involved in TCR cell signaling, such as Ptpn6 Y564, Ptpn12 S673, Lck S166, Grap2 T254, Lat T62 and Y67, and Map3k8 S141. Several of these phosphorylation sites have already been studied in the context of T cell exhaustion. Notably, Grap2 T254 (T262 in humans) is increased in T_EX_, consistent with its function negatively regulating T cell activation. As an adaptor protein, Grap2 plays a pivotal role in linking the TCR to downstream signaling cascades, thereby mediating T cell activation and effector function^66–68^.

Leveraging the longitudinal nature of our data, we examined PTM regulation on key proteins (Fig.4H). PD1 Y225 and Y248^15, 69^ – located within the ITIM and ITSM motifs, respectively – exhibited contrasting phosphorylation patterns in T_EX_ vs T_EFF_/T_MEM_, with Y225 downregulated and Y248 upregulated in T_EX_ (Fig.4G, H). This is consistent with functions of these PD1 phosphosites: p-Y248 is known to mediate inhibitory signaling via SHP-2 and SHIP recruitment, dampening T cell activation^70^, while Y225 (Y223 in humans) is dispensable for SHP-2 binding and downstream PD1 signaling in T cells^15^. On SHP-1/Ptpn6, a negative regulator of TCR signaling, we also observed a significant increase in Y564 phosphorylation in T_EX_. This is consistent with published work, indicating phosphorylation of Y536 or Y564 on SHP-1/Ptpn6 relieves auto-inhibition, allowing its phosphatase activity to dampen signaling^71–74^. We also observed upregulation of Lag3 S476, embedded within its FSALE motif (conserved between mouse 475-479 and human 481-487), a sequence essential for LAG3’s suppressive activity. Lag3 S476 (S484 in humans) is dispensable for Lag3 inhibitory activity, however considering its conservation (see Fig.5D) in mouse and humans, it may have another uncharacterized regulatory function^75, 76^.

**Figure 5:**
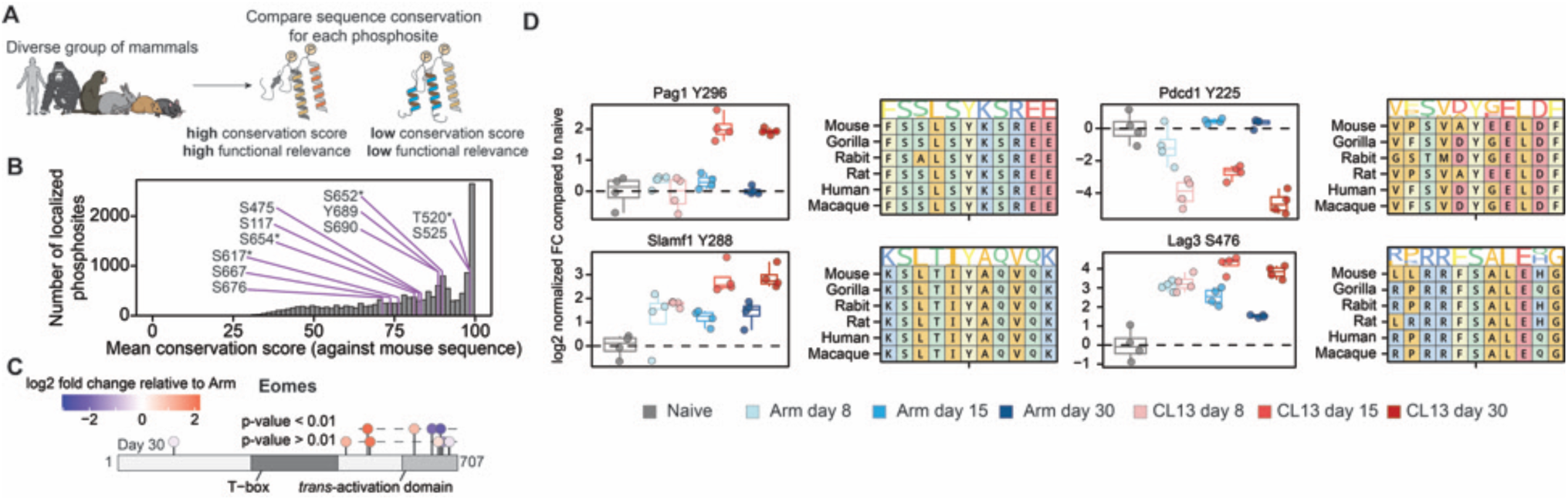
Phosphoproteomic profiling of T cells identifies regulatory exhaustion-specific modification sites conserved across mammalian species. (**A**) Comparative analysis of phosphorylation site conservation across multiple mammalian species. (**B**) Histogram quantifying phosphosite conservation scores across mammalian species compared with mouse. Sites on Eomes are highlighted in purple; asterisks (*) denote Eomes sites with significant phosphorylation difference between T_EX_ and T_MEM_ at day 30. (**C**) Schematic indicating quantified Eomes phosphorylation sites compared between T_EX_ and T_MEM_ (Arm) at day 30. (**D**) Boxplots display phosphorylation levels at individual phosphosites along the T cell differentiation trajectory. Amino acid sequence alignments show protein conservation across six mammalian species, with the phosphorylated residue centered and marked by a tick. N = 4 for all proteomics samples.

To prioritize functionally relevant phosphorylation sites for potential pharmacological interventions, we assessed sequence conservation of phosphosites quantified in our mouse dataset across five other mammalian species (human, gorilla, macaque, rabbit, and rat), based on the premise that conserved sites are more likely to have functional significance (Fig.5A-B). We quantified conservation of phosphosites on a scale of 0-100, with 100 representing perfect conservation across all six mammalian proteomes (Fig.5B, Supplementary Table 6). As an example, we studied conservation of phosphorylation sites on Eomes, a transcription factor associated with T cell exhaustion (Fig.5B-C). We observe a conservation score range of 71-98 across all phosphorylation sites in Eomes, and a range of 76-98 for four significantly altered sites (T_EX_ vs T_MEM_ on d30). Among these four significant sites, T520 and S617 are upregulated, and S652 and S654 are downregulated in T_EX_ (Fig.5C). The functions of these phosphosites are unknown, but three of the four were located within the transactivation domain which is responsible for transcriptional activity^77^, and two (S652 and S654) are located adjacent to a putative nuclear localization signal located at residues 662-665^78^, with potential relevance to the nuclear trafficking of Eomes.

We also examined other phosphosites of interest with high conservation (Fig.5D, Supplementary Tables 6 and 7). For example, we observe increased phosphorylation of Pag1 at Y296 in T_EX_. Phosphorylated Pag1 (also known as Cbp) regulates T cell signaling by recruiting Csk to the membrane, where Csk attenuates TCR signaling by negatively regulating Src kinases^79, 80^. Importantly, mutational studies demonstrate that Y296 phosphorylation is crucial for Csk binding: substitution of Pag1 Y296 with phenylalanine weakens the Csk interaction, while combined mutation of Y296 and Y314 abolishes it entirely. This evidence links elevated Pag1 Y296 phosphorylation in T_EX_ to Csk inhibition of TCR signaling^81^. We also observed increased Slamf1 p-Y288 (Y281 in humans) in T_EX_. This residue is located within the ITSM motif of Slamf1/CD150, a T-cell co-stimulatory molecule, and Slamf1 Y288 phosphorylation mediates binding to SHP-2 and SHIP, two key negative regulators of T cell signaling^70^. Finally, we also highlight high conservation of PD1 Y225 and Lag3 S476 (discussed earlier).

Our analysis identifies known and novel phosphorylation modifications in exhausted T cells, suggesting potential mechanisms of immune regulation meriting further investigation.

## Discussion

This study presents the first in vivo longitudinal proteomic and phosphoproteomic profiling of antigen-specific CD8 T cell differentiation during both chronic and acute viral infections. By contrasting T cells from chronic (T_EX_) and acute (T_EFF_/T_MEM_) infection contexts, we identified differentially abundant proteins and phosphosites that distinguish these states.

Most systems-level T cell exhaustion analyses have been performed using NGS-based technologies. While transcriptomics captures a momentary view of RNA expression and epigenetics sheds light on gene regulation, these techniques do not directly measure the global proteome, which comprises the molecular machines and functional components that orchestrate cellular functions. Reassuringly, we observe a correlation between transcriptomic and protein abundance data, and thus many of the top exhaustion-associated hits in our protein abundance dataset have previously been identified and validated^8, 21^. These include classical inhibitory receptors: PD1/*Pdcd1*, Lag3, TIM3/*Havcr2*, 2B4/*Cd244*, and TIGIT as well as transcription factors implicated in exhaustion: Tox and Eomes. In addition, our protein abundance analysis identified several proteins elevated in T_EX_ that were not highlighted by sequencing-based analyses. This discrepancy is explained by limited correlation between protein abundance and mRNA levels, also observed in diverse contexts^10, 19, 45^, as well as limited read depth in scRNAseq analyses, which we’ve observed for some exhaustion-associated transcripts (see Fig.2G). Based on current knowledge of these gene functions, they warrant further analysis towards their role in T cell function.

Our phosphorylation analysis highlighted state-specific PTMs modulating immune signaling, representing opportunities for novel insights into T cell regulation and immunotherpeutic strategies. While a few phosphosites in our dataset have been implicated in key T cell signaling pathways, e.g., PD1 Y248^15, 69^ and Grap2 T254^66–68^, the majority of phosphosites identified from our phosphoproteomics dataset are not implicated in T cell regulation, and some are entirely novel. Interestingly, our data indicate that Eomes is rich in phosphosites that are state-dependent, reflecting a previously unrecognized layer of regulation of the T cell differentiation program. Eomes is implicated in both the memory state and the late exhaustion state, and displays differentially regulated phosphosites that are enriched in either state, suggesting a potential involvement of these PTMs in regulating the differential role of the same transcription factor under various conditions. Thus, beyond known regulatory sites, our dataset highlights novel phosphosites with potential functional relevance, providing new insights into T cell differentiation. More broadly, biochemical properties such as post-translational modifications (PTMs) are not accessible by NGS-based readouts, and these represent new opportunities for studying immune regulatory biochemistry and identifying therapeutic strategies.

Moving forward, a key challenge in PTM analysis is the functional annotation of modification sites. For example, large-scale MS-based studies have reported over 200,000 phosphorylation sites in human proteins, yet their functional significance remains largely unknown^85^. Recent advances in CRISPR/Cas9 have yielded efficient base editors applicable to unbiased high-throughput mutational analyses of protein residues^86, 87^, and new innovations continue to expand the precision and efficiency of gene editing^88^. Utilizing the phosphoproteomic data in this study to design targeted mutational libraries, future work will apply precision gene editing to identify the phosphosites and protein surfaces mediating T cell differentiation and exhaustion. Phosphorylation is only one of hundreds of PTMs, and further proteomic analyses will explore other biochemical modifications mediating T cell differentiation and exhaustion.

In addition to identifying individual phosphosites, a higher-order interpretation of our phosphoproteomics data is provided by kinase substrate enrichment analysis (KSEA), which infers state-specific kinase activity based on phosphorylation sites. We validated our approach by analyzing the activity of CDKs associated with cell proliferation, which reflected expected activity in both acute and chronic settings^23^. Additionally, KSEA highlighted several exhaustion-associated kinases, several of which have been functionally validated in previous work. Crucially, there are over 80 FDA-approved kinase inhibitors and approximately 180 orally effective kinase inhibitors in clinical trials^89^, providing opportunities to leverage our data towards clinical translation.

Finally, our analyses of the evolutionary conservation of phosphosite motifs across mammalian species further reinforce the value of our dataset. Mouse models, especially LCMV, have been invaluable in dissecting the T_EX_ biology that was translated into human settings. Since the inception of T cell exhaustion studies, a plethora of work has confirmed the conserved T_EX_ core program across species (mice/macaques/humans) and variable chronic settings^90–94^. Given the conservation of T cell regulatory mechanisms across vertebrates, our analysis highlights high priority phosphosites to guide the field’s focus.

A key strength of the LCMV system is the differential analysis of chronic versus acute infection, permitting isolation of specific molecular features differentiating T_EX_ from T_EFF_/T_MEM_. For brevity, we chose to focus largely on this analysis to highlight the most impactful and translatable features in our dataset. Isolated trajectories of naïve to T_EX_, or naïve to T_EFF_/T_MEM_, are also provided in our extended data, and can be leveraged to identify and study novel biochemical mechanisms underlying T cell differentiation.

The fields of immunology and cancer research are on a quest for novel interventions to enhance the efficacy and durability of immunotherapy^95^. One understudied aspect of T cell biology is profiling their biochemical dynamics. Here, our study provides the first-of-its-kind dataset dissecting the kinetics of protein abundance and phosphorylation in antigen-specific T cells. Our protein abundance profiling of T_EX_ revealed novel proteins with potential roles in exhaustion biology not previously highlighted by transcriptional readouts, highlighting the importance of proteomic analysis to identify the ‘ground truth’ of protein expression. Furthermore, investigation of PTMs provides a biochemical perspective that is largely inaccessible through sequencing-based approaches. This comprehensive biochemical data is essential for dissecting the regulatory networks that govern T cell exhaustion programs and fate decisions, and ultimately developing therapeutic strategies to modulate immunity and combat T cell dysfunction.

<u>NOTE</u>: In-depth validation of key findings from our proteomics-guided analysis is currently underway. This includes confirming the roles of newly identified proteins and phosphorylation sites in T cell signaling and exhaustion. These follow-up studies will be essential for translating our large-scale proteomic observations into mechanistic insights and potential therapeutic strategies.

## Acknowledgments

This work was supported by an Emory MP3 Seed Grant (121861), a Winship Cancer Center Invest$ Pilot Award (147150), and Department of Pathology and Laboratory Medicine Start-up funds to MSA and DEG. CMB was supported by a Swedish Research Council grant (2023-00510). ASF was supported by the National Institutes of Health through the Initiative for Maximizing Student Development (IMSD) T32 Fellowship (T32GM148391-01). The authors would like to thank members of the Hakeem and Gordon labs, as well as members of the Pathology Advanced Translational Research Unit (PATRU) and Emory Vaccine Center (EVC) for their valuable discussions and comments. We would also like to thank the Emory Flow Cytometry Core (EFCC) and the Emory Integrated Genomics Core (EIGC) for their support. This study was supported in part by the Emory Flow Cytometry Core (EFCC), one of the Emory Integrated Core Facilities (EICF), and is subsidized by the Emory University School of Medicine. Additional support was provided by the National Center for Georgia Clinical & Translational Science Alliance of the National Institutes of Health under Award Number UL1TR002378. Research reported in this publication was supported in part by the Emory Integrated Genomics Core (EIGC) shared resource of Winship Cancer Institute of Emory University and NIH/NCI under award number P30CA138292. We thank the NIH Tetramer Core Facility (contract number 75N93020D00005) for providing tetramers used in this study (H2-Db LCMV-GP33-41, KAVYNFATC). The NIH Tetramer Core Facility is supported through National Institute of Allergy and Infectious Diseases (NIAID) and co-funding from the National Cancer Institute (NCI). The content is solely the responsibility of the authors and does not necessarily represent the official views of the National Institutes of Health.

---

**Extended Data Figure 1:**
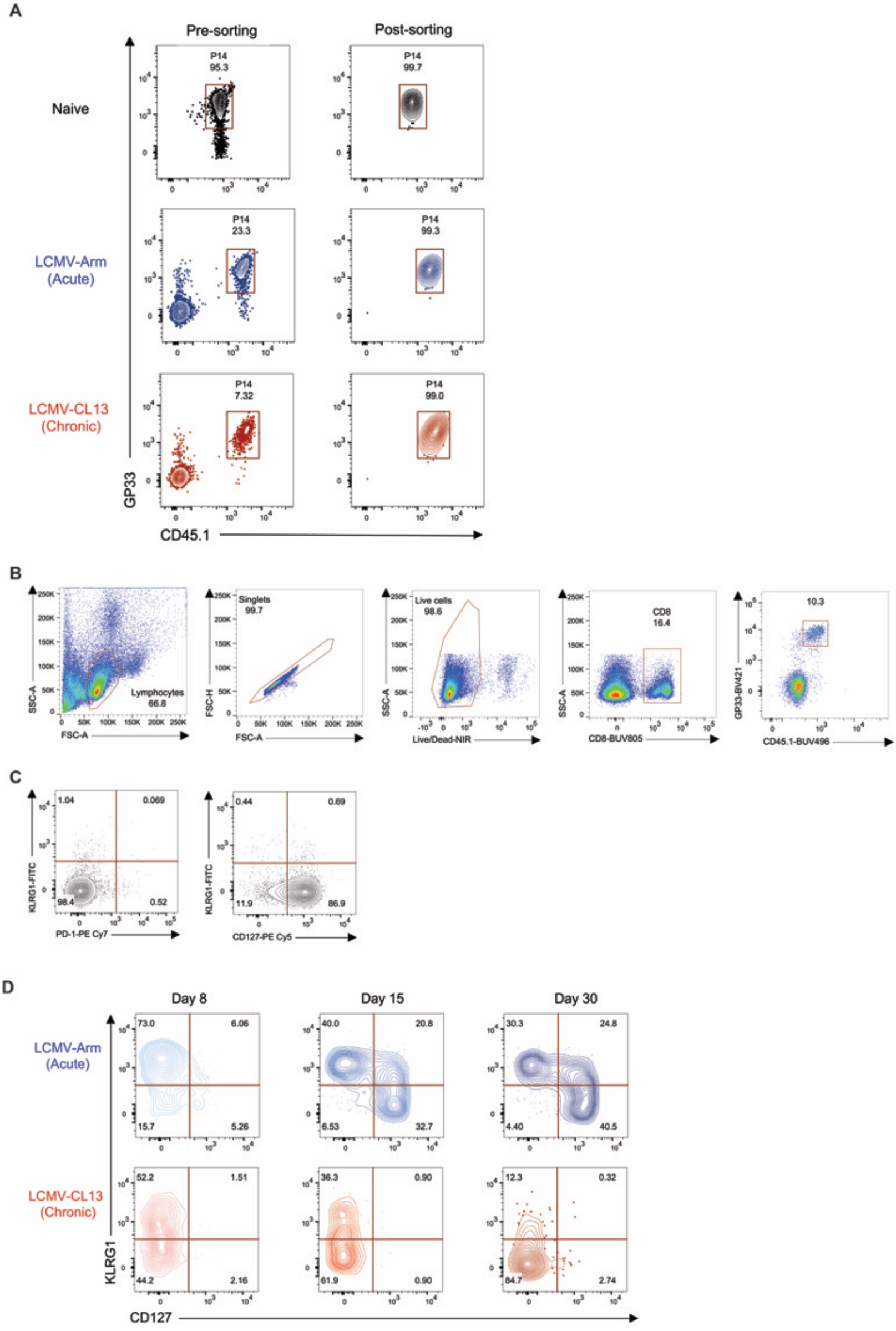
Flow cytometry validation of the phenotype and sorting purity of antigen-specific CD8 T cells used for proteomic profiling by mass spectrometry-based proteomics. (**A**) Representative dot plots reflecting the sorting purity (post-sorting, right panels) compared to the original frequencies pre-sorting. (**B**) Gating strategy for flow cytometry analysis. Representative dot plots of the phenotypic analysis of for the markers KLRG1, CD127, and PD-1 for naïve CD8 T cells (**C**), and for the markers KLRG1 and CD127 for P14 cells longitudinally during LCMV-Arm and LCMV-CL13 infection (**D**).

**Extended Data Figure 2:**
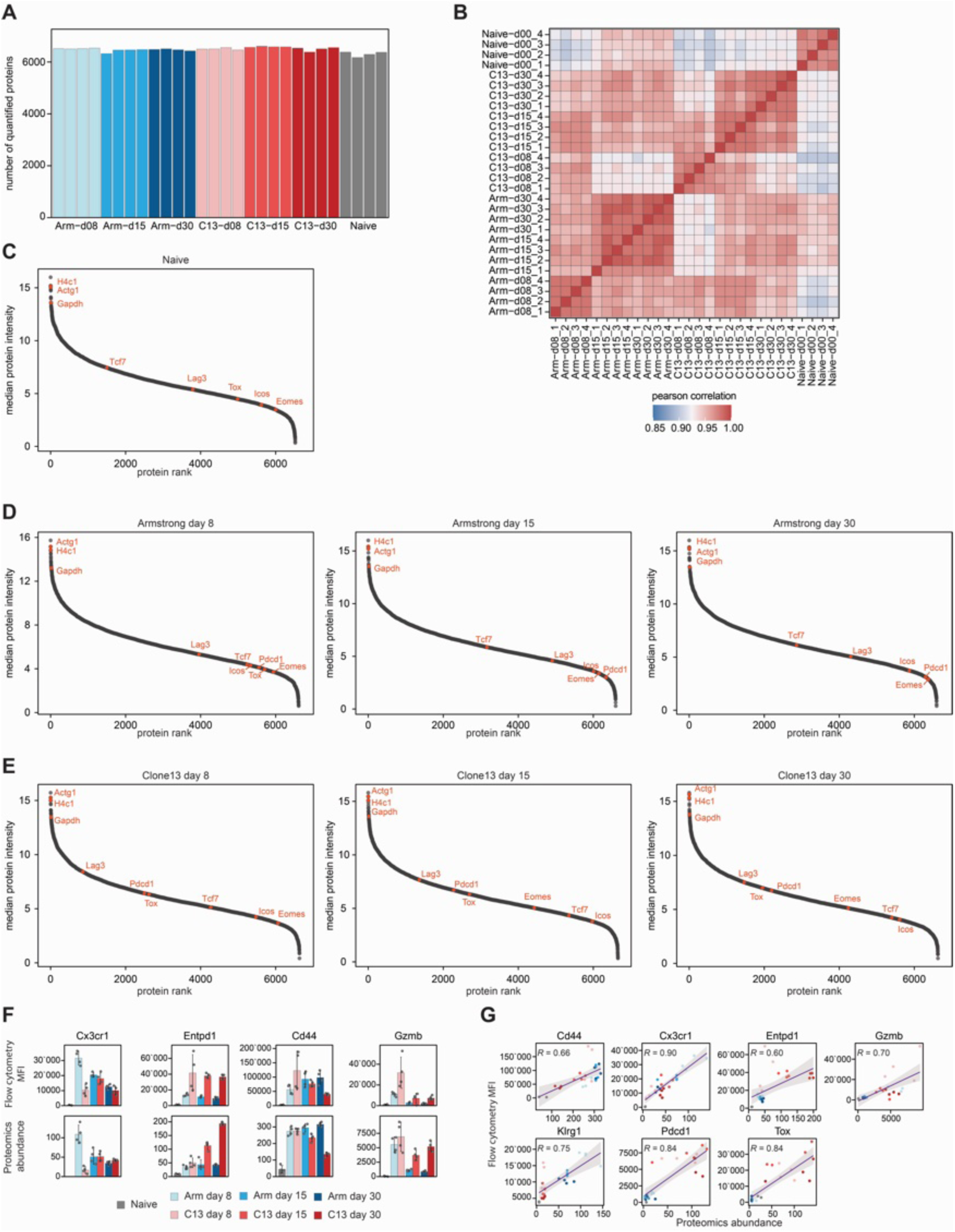
Assessment of protein abundance profiling of antigen-specific CD8 T cells in the context of acute and chronic infection. (**A**) Number of quantified proteins in each proteomics sample. (**B**) Correlation between the different samples from all four biological replicates. (**C-E**) Dynamic range for the protein abundance quantification for the naive (**C**), acute (**D**), and chronic (**E**) samples. (**F**) Comparison of protein expression assessed through flow cytometry (top) and proteomics (bottom). The absence of a measurement bar indicates no quantification. (**G**) Pearson correlation between protein expression levels assessed through flow cytometry (Y-axis) and proteomics (X-axis). Naive B6 mice were used as staining controls in (**F-G**).

**Extended Data Figure 3:**
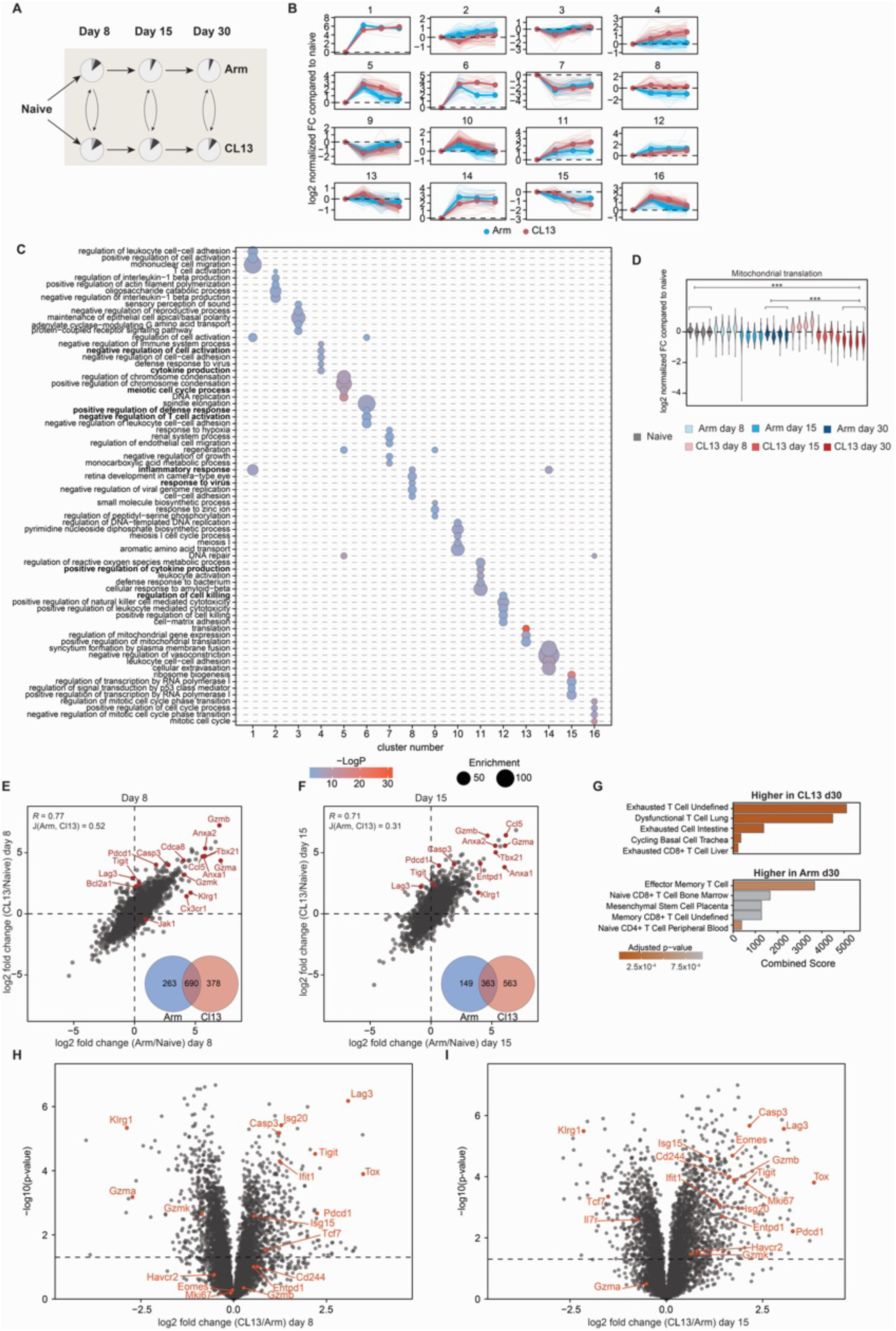
Protein alterations in the context of T cell exhaustion compared to differentiation in an acute context. (**A**) Schematic representation of analyzed samples used in an unsupervised clustering approach to identify protein expression patterns specific to exhausted T cells. (**B**) Longitudinal protein expression dynamics across all clusters from Figure 3A, with bold lines indicating the average protein trajectory over time. (**C**) Gene Ontology (GO) enrichment analysis of protein clusters identified in Figure 3A, highlighting functional pathways associated with exhaustion. (**D**) Distribution of protein expression changes for mitochondrial translation components across all samples, illustrating metabolic and cell cycle regulatory shifts. (**E, F**) Scatter plots comparing protein expression changes in acute versus chronically infected T cells relative to naïve cells at days 8 and 15 post-infection. The Venn diagram quantifies the overlap of significantly regulated proteins between the two conditions. Pearson correlation was used to assess sample similarity, while the Jaccard index evaluated the overlap of significantly regulated proteins. (G) Signatures of differentially regulated proteins from Figure 3D, identified using the CellMarker database^2^. (**H, I**) Volcano plots illustrating differential protein abundance in CD8⁺ T cells from Armstrong-versus Clone 13-infected mice at days 8 and 15 post-infection. P-values were calculated using a two-sided Student’s t-test with equal variance. N = 4 for all proteomics samples.

**Extended Data Figure 4:**
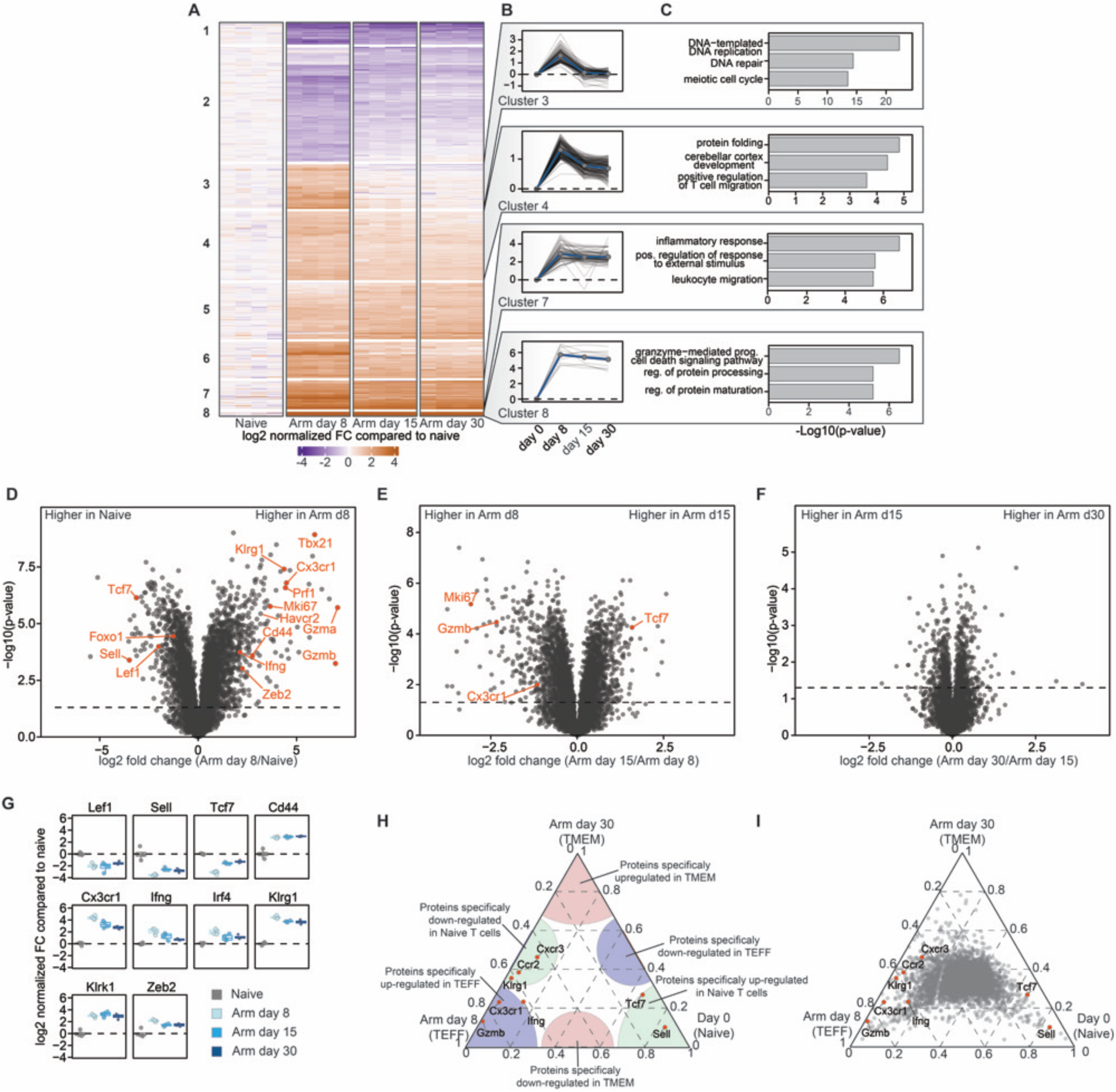
Expression profiles of proteins defining T cell differentiation during acute LCMV-Arm infection. (**A**) Unsupervised hierarchical clustering of differentially expressed proteins throughout the LCMV Armstrong (acute) infection time course. (**B**) Longitudinal expression dynamics of proteins within selected clusters. The bold blue line represents the average protein change. (**C**) Gene Ontology (GO) enrichment analysis of selected protein clusters, highlighting biological processes associated with different phases of T cell differentiation. (**D-F**) Volcano plot depicting differentially expressed proteins in naïve CD8⁺ T cells compared to antigen-experienced effector CD8⁺ T cells at day 8 post-infection, d8 (left), d15 (middle, and d30 (right). P-values were calculated using a two-sided Student’s t-test with equal variance. (**G**) Longitudinal expression trends of selected proteins across the differentiation trajectory of CD8⁺ T cells, tracking key regulators and markers from naïve to effector and memory states. (H) Explanatory ternary plot. (**I**) Ternary plot depicting protein-specific expression patterns across the transition trajectory from naïve, to effector (Arm d8), then memory (Arm d30) T cells, illustrating key regulators of T cell differentiation. Named proteins act as reference points. N = 4 for all proteomics samples

**Extended Data Figure 5:**
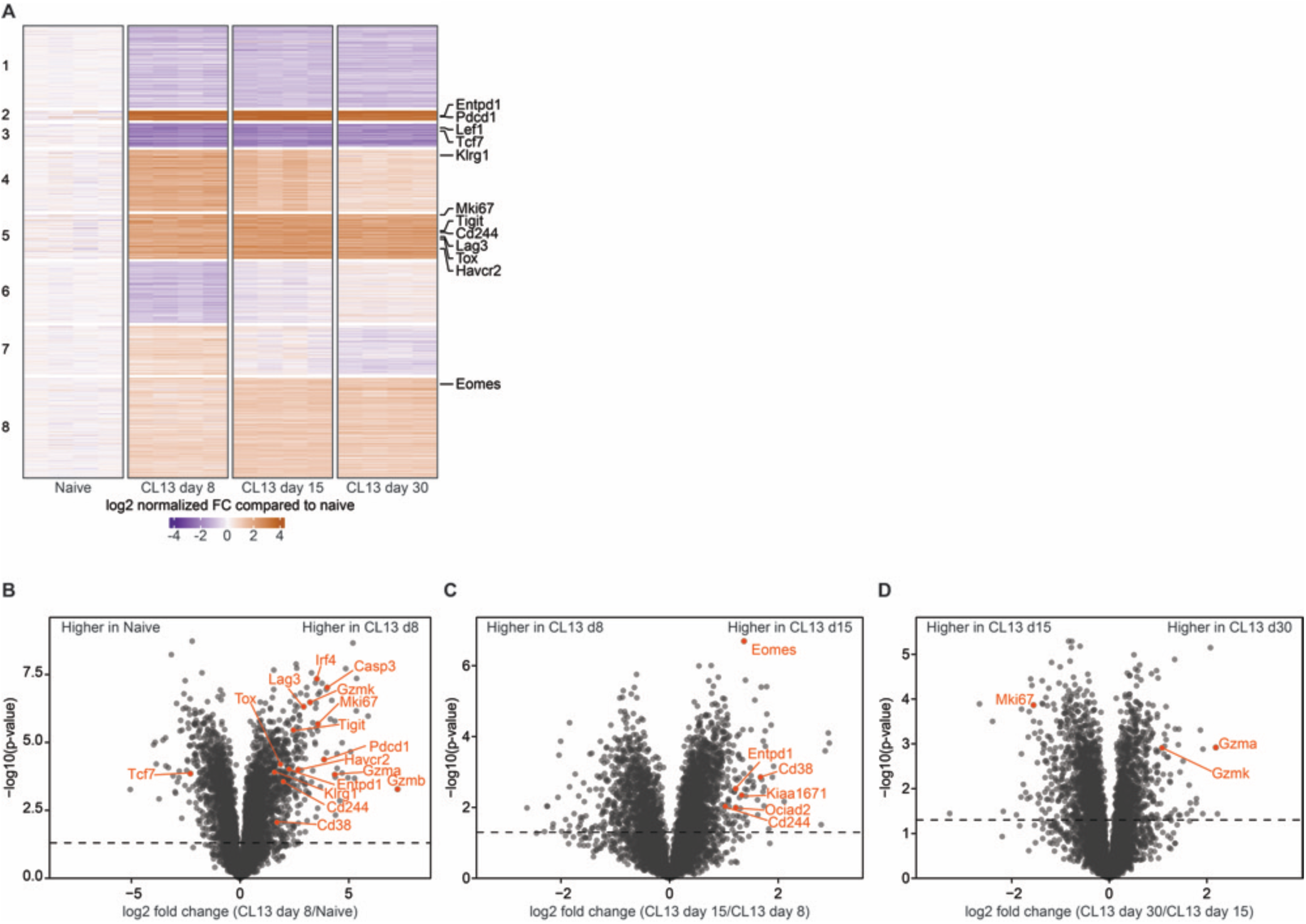
Expression profiles of proteins defining T cell exhaustion. (**A**) Unsupervised hierarchical clustering of differentially expressed proteins throughout the LCMV-CL13 (chronic) infection time course. (**B-D**) Volcano plots depicting differentially expressed proteins in naïve versus early-Tex d8 (left), d8 versus d15 (middle), and d15 versus d30 (right). P-values were calculated using a two-sided Student’s t-test with equal variance. N = 4 for all proteomics samples.

**Extended Data Figure 6:**
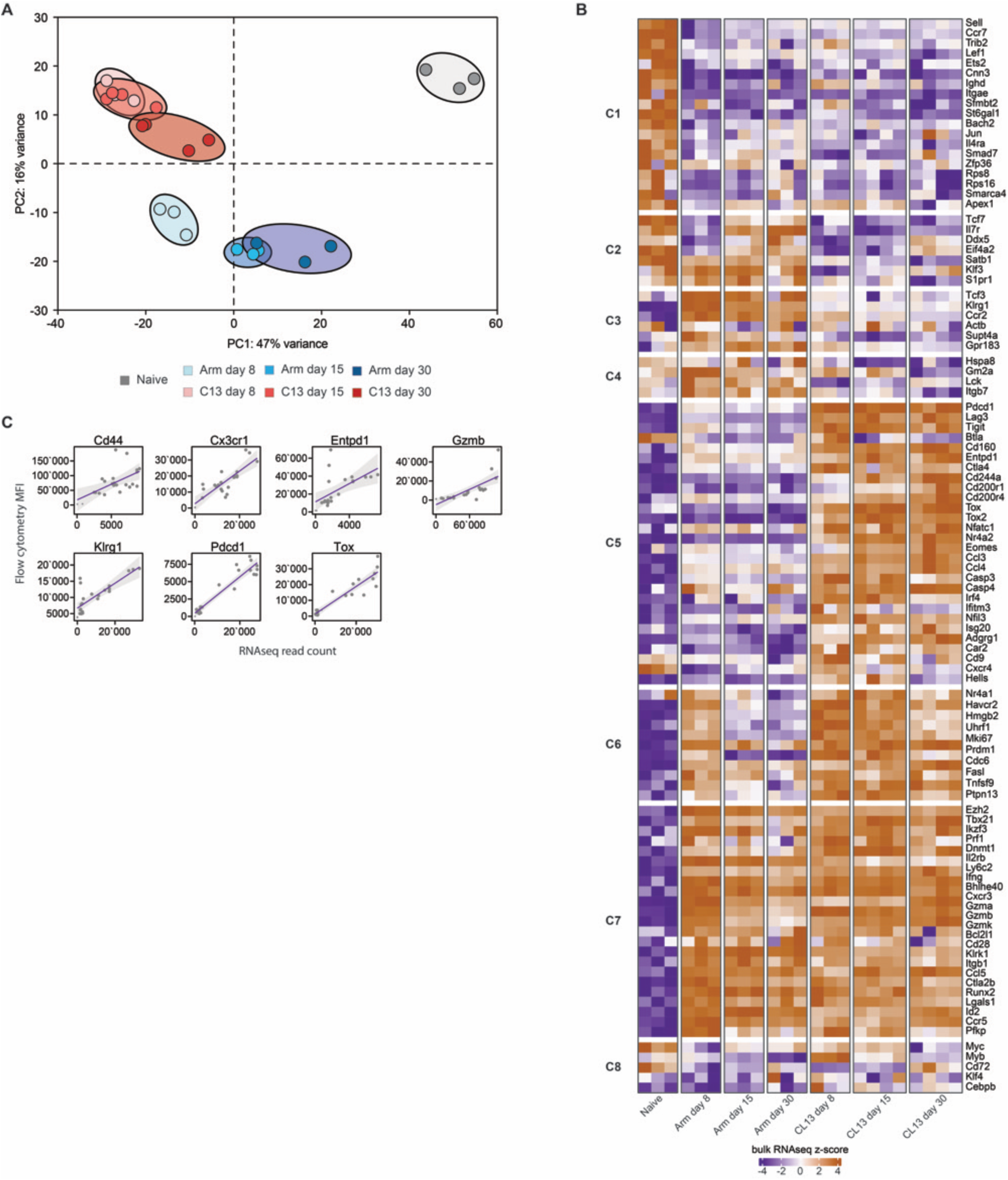
Transcriptomic analysis and correlation of changes in transcript abundance compared with protein expression. (**A**) Principal component analysis (PCA) of RNA sequencing samples. (**B**) Differentially expressed genes in the different samples, reflecting previously reported state-specific modules^3, 4^. Each column represents a biological replicate. (**C**) Pearson correlation analysis comparing protein expression levels measured by flow cytometry (Y-axis) with RNA sequencing counts (X-axis). Naïve B6 mice were used as staining controls. N = 3-4 for RNA sequencing.

**Extended Data Figure 7:**
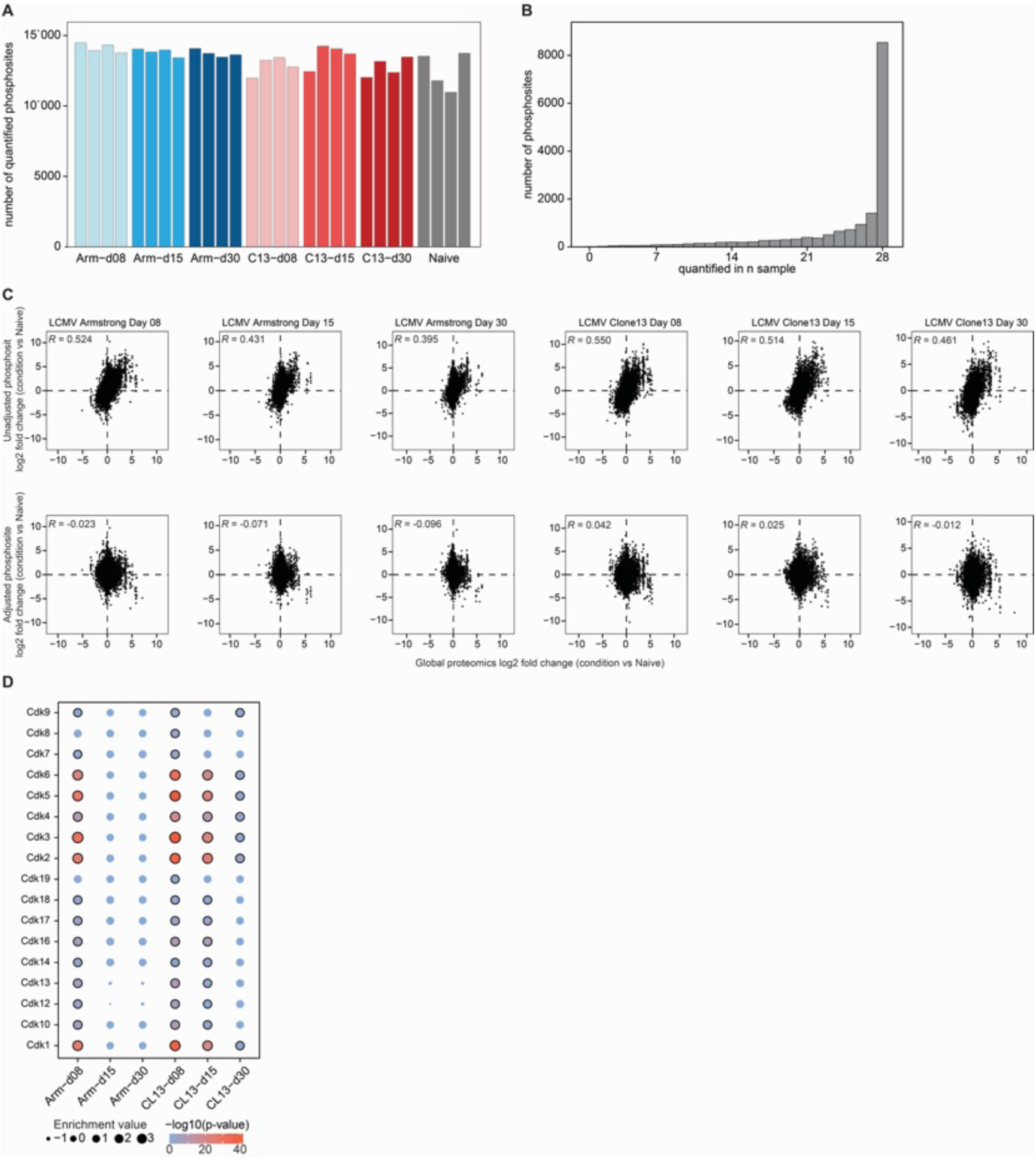
Phosphoproteomic analysis and kinase activity in T cell differentiation. (**A**) Quantification of phosphosites in each sample, showing the number of identified sites per condition. (**B**) Overlap of phosphosite quantification across all samples, highlighting shared and unique phosphosites. (**C**) Correlation analysis between protein expression measurements and phosphosite changes, both before and after adjusting for intrinsic protein expression changes during T cell differentiation. Sample correlation was determined using Pearson correlation, with only one replicate shown per sample. (**D**) Kinase-substrate enrichment analysis (KSEA) for all cyclin-dependent kinases (CDKs), comparing each time point to naïve T cells.

